# Pangenome reconstruction in rats enhances genotype-phenotype mapping and novel variant discovery

**DOI:** 10.1101/2024.01.10.575041

**Authors:** Flavia Villani, Andrea Guarracino, Rachel R Ward, Tomomi Green, Madeleine Emms, Michal Pravenec, Pjotr Prins, Erik Garrison, Robert W. Williams, Hao Chen, Vincenza Colonna

## Abstract

The HXB/BXH family of recombinant inbred rat strains is a unique genetic resource that has been extensively phenotyped over 25 years, resulting in a vast dataset of quantitative molecular and physiological phenotypes. We built a pangenome graph from 10x Genomics Linked-Read data for 31 recombinant inbred rats to study genetic variation and association mapping. The pangenome includes 0.2Gb of sequence that is not present the reference mRatBN7.2, confirming the capture of substantial additional variation. We validated variants in challenging regions, including complex structural variants resolving into multiple haplotypes. Phenome-wide association analysis of validated SNPs uncovered variants associated with glucose/insulin levels and hippocampal gene expression. We propose an interaction between *Pirl1l1*, chromogranin expression, *TNF-α* levels, and insulin regulation. This study demonstrates the utility of linked-read pangenomes for comprehensive variant detection and mapping phenotypic diversity in a widely used rat genetic reference panel.

## Introduction

*Rattus norvegicus* has a long history of scientific research and has been used as a model organism to study various human diseases.^1^ One notable example is the HXB/BXH (referred to as HXB in general hereafter) family of recombinant inbred (RI) strains of rats. The HXB family started with reciprocal crosses between the SHR/OlaIpcv (H) and BN-Lx/Cub (B) strains, with the goal of creating distinct genetic lines. Currently, there are a total of 32 RI strains, all having undergone more than 50 generations of inbreeding. Among them, 21 belong to the HXB/Ipcv group, and 11 to the BXH/Cub group, with DNA samples preserved from almost all the original strains for the purpose of constructing genetic maps.^2^

The HXB is currently the most widely used RI family of laboratory rats and therefore a great resource for the identification and mapping of quantitative trait loci (QTL) associated with complex phenotypes, motivating efforts to produce a comprehensive catalog of genetic variants. HXB have been utilized for studying the genetics of a wide range of phenotypes related to human diseases, such as blood pressure,^3^ cardiac conduction,^4^ Pavlovian conditioning,^5^ ethanol metabolism^6^ and hippocampal neurogenesis.^4^

Pangenomic methods directly compare all genomes to each other, enabling a comprehensive analysis of genomic diversity and relationships.^7^ These methods have previously proven successful in representing human genetic variation^8^ and addressing long-standing evolutionary questions.^9^ The design, implementation, and application of pangenomic models constitute an active area of research. Ideally, pangenomes are constructed from long read data, but this technology is not always available, and the accuracy and utility of pangenome construction from suboptimal sequence data have been insufficiently explored.

In this study, we build a pangenome graph from 10x Chromium Linked-Reads sequence data from 31 HXB rats plus BN/NHsdMcwi (mRatBN7.2 reference genome), and use newly identified/discovered genetic variation to perform phenome-wide association analysis and explore complex genetic variability within structural variants. We demonstrate the impact of using pangenomes in the mapping of complex traits through examples of genetic association of novel variants discovered from the pangenome and traits related to insulin and glucose metabolism.

## Results

### Pangenome building from Linked-Reads

We performed whole genome sequencing using the 10x Chromium Linked-Read technology (average coverage depth 109) and *de novo* genome assembly of 31 inbred strains from the RI rat panel^10^ **(Figure S1A)**. Evaluation of genome assembly within genic regions^13^ shows that the majority of the genes are complete and present in a single-copy configuration **(Figure S1)**, which is indicative of a high quality.

Using classical reference-based approach as implemented in joint-calling (JC) of variants using DeepVariant/GLnexus,^14^ we identified 7,520,223 variant sites from the HXB/BXH panel, approximately 24.1 % – 53.5 % of the alleles at these sites of each RI strain were derived from SHR/OlaIpcv. We found that the majority of the strains were highly inbred, with close to 98% of the variants being homozygous, one exception was BXH2, in which 7.7% of variants were heterozygous, likely due to a recent breeding error.^6^ Based on these results, we considered the assembly as haploid and applied PGGB^8^ to build pangenome graphs for all chromosomes from *de novo* genome assemblies of the full HXB family (generated from 10x Linked-Reads) and the mRatBN7.2 (rn7) reference genome. The lengths of such chromosome pangenome graphs were calculated as the sum of their node lengths and after removing uncertain sequence stretches they were, on average, 1.05 times greater than the corresponding chromosome length of the reference mRatBN7.2, suggesting that the pangenome captures genetic variation beyond that contained in the reference **(<u>Table S1</u>)**. This is expected because the pangenome graphs we built represent the genetic diversity of all the strains they represent, including Single Nucleotide Polymorphisms (SNPs), Insertions-deletions (Indels), and Structural Variations (SVs). In particular, each allele would be represented with a dedicated node, therefore increasing the total graph length. For this reason, pangenome graphs can be longer than the individual genomes they represent. To demonstrate the utility of a pangenomic approach, we focused on chromosome 12 (chr12), one of the shortest chromosomes. The pangenome graph of scaffolds that map to chr12 consists of 1M nodes, 1.6M edges, and 28.5k paths, with a total length of 78M bp **(<u>Table S2</u>, Figure 1A**).

**Figure 1.**
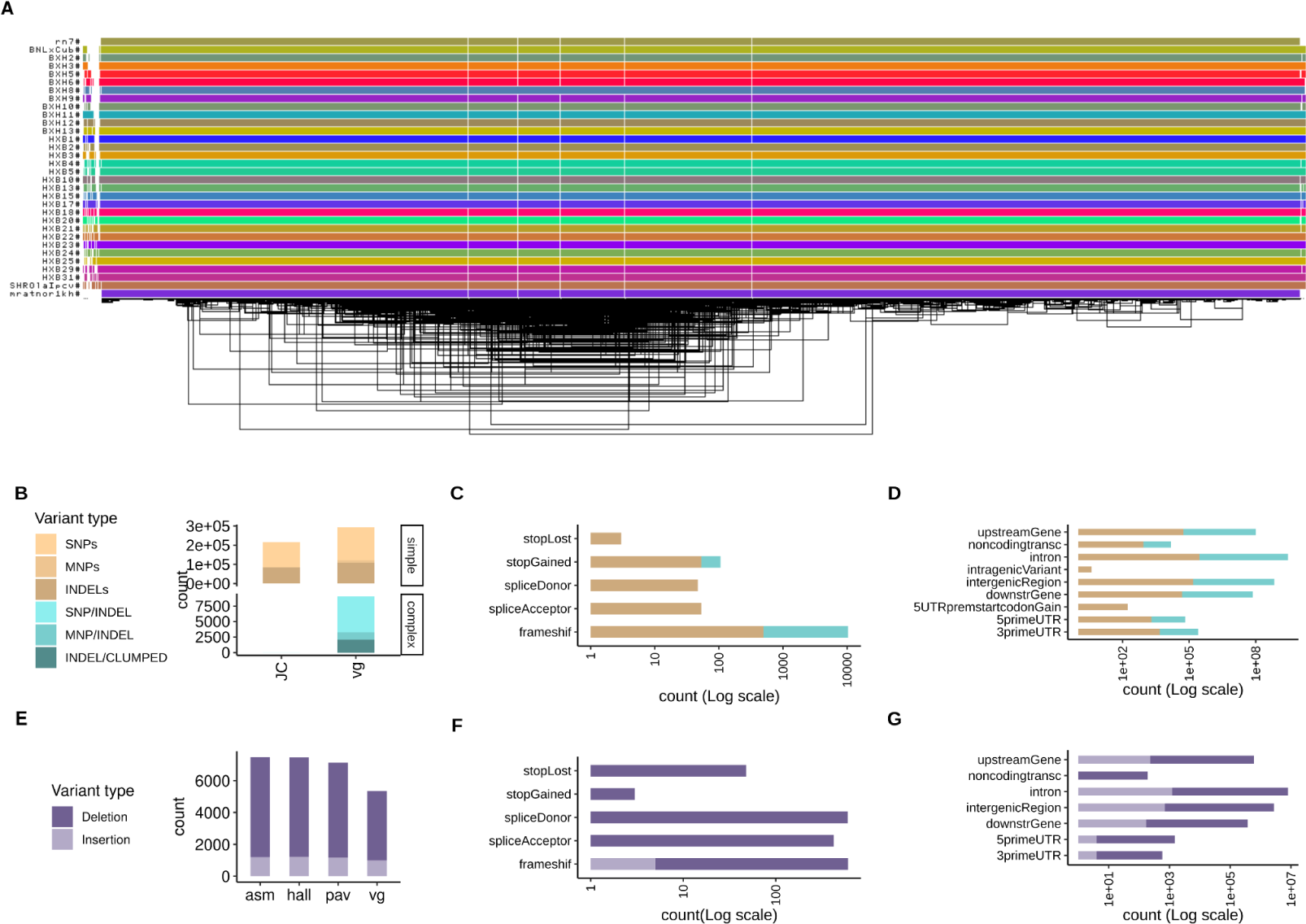
Pangenome graph of chromosome 12 from 31 HXB/BXH rats and genetic variants derived from it. **(A)** odgi-viz visualization of the pangenome graph for chromosome 12 from 31 rats. Horizontal colored lines denote haplotypes, with vertical white stripes representing insertions/deletions. Includes the mRatBN7.2 reference genome (top line). **(B)** Comparison between small variants (<50bp) called from the pangenome graph using (vg) with variants joint-calling directly from 10x data using GLnexus (JC), stratified by genome complexity. vg called 36% and 99.95% more simple and complex variants, respectively, compared to JC. **(C-D)** Predicted functional annotations for small variants called from the pangenome graph. **(E)** Comparison of three assembly-based methods (asm, hall, pav) and one graph-based method (vg) to call structural variants (SVs, >50bp) **(F-G)** Predicted functional annotations for SVs called from the pangenome graph.

### Small variants analysis and validation in the pangenome

Small variants (<50 bp in length) were called from the pangenome using graph decomposition to identify bubbles corresponding to non-overlapping variant sites, as implemented in the vg toolkit.^15^ We also performed a classical reference-based approach as implemented in joint-calling (JC) of variants using DeepVariant/GLnexus.^14^ Variant calling from the pangenome of chr12 identified 302,048 variants (SNPs, multi-nucleotide polymorphisms (MNPs), Indels, and complex, i.e. nested mixed variants) *versus* 215,431 variants called by JC. Of these, vg called 36% and 99.95% more simple and complex variants respectively (**Figure 1B**). With a focus on size up to 50 bp (small variants), we predicted functional consequences of variants on genes and proteins **(<u>Table S3</u>)**. The majority of variants were classified as modifiers (99.0%). A smaller proportion of variants had a high impact (0.1%) and were located within 364 protein-coding genes. For each class of variant impact (high, moderate, low, and modifier) annotations were more abundant for simple variants (**Figure 1C-D**). Finally, with respect to gene structure, the majority of variants were located in intergenic regions and introns **(<u>Table S3</u>)**.

Small variants called from the pangenome using vg were validated over the JC call set (gold standard) in the SHR/OlaIpcv sample **(<u>Table S4</u>)**, which represents one of the best assemblies among the strains in our pangenome in terms of quality **(<u>Table S5</u>, Figure S1B)**. Genome-wide accuracy of vg calls was ∼90% **(Figure S2)**, with the exceptions of chromosome Y and 16, possibly due to the complexities in assembling chromosome Y and the abundance of complex variants for chromosome 16 **(Figure S2C)**. Further validation by resequencing was restricted to twenty SNPs within easy regions^16^ of chr12. Selection criteria for these small variants were: support by original reads determined by mapping reads against the pangenome with vg giraffe,^17^ and location in regions supposed to be particularly challenging (14 were within repetitive regions and 6 within 1500 bp from repetitive regions). Twelve out of the 20 variants were confirmed by Sanger sequencing **(<u>Table S6</u>)**, which translates into a 0.6 rate of validation. Approximately 90% of variants (7 out of 8) that failed the validation were within repeats.

### Novel pangenome variants associated with glucose, insulin and chromogranin expression

We leveraged phenotypic data present in GeneNetwork^18^ to perform phenome-wide association analysis (PheWAS) for the twelve SNPs validated through Sanger sequencing. PheWAS was restricted to 30 rat strains with phenotypic data, and 60 phenotype classes with at least two phenotypes per class. Two SNPs were significantly associated with six phenotypes within five phenotype classes (LRS>16, **Figure 2A-B, <u>Table S7</u>**). We further considered three of the six significant associations that were validated using a linear mixed model corrected for kinship implemented in GeneNetwork **(Figure 2C-E)**.

**Figure 2.**
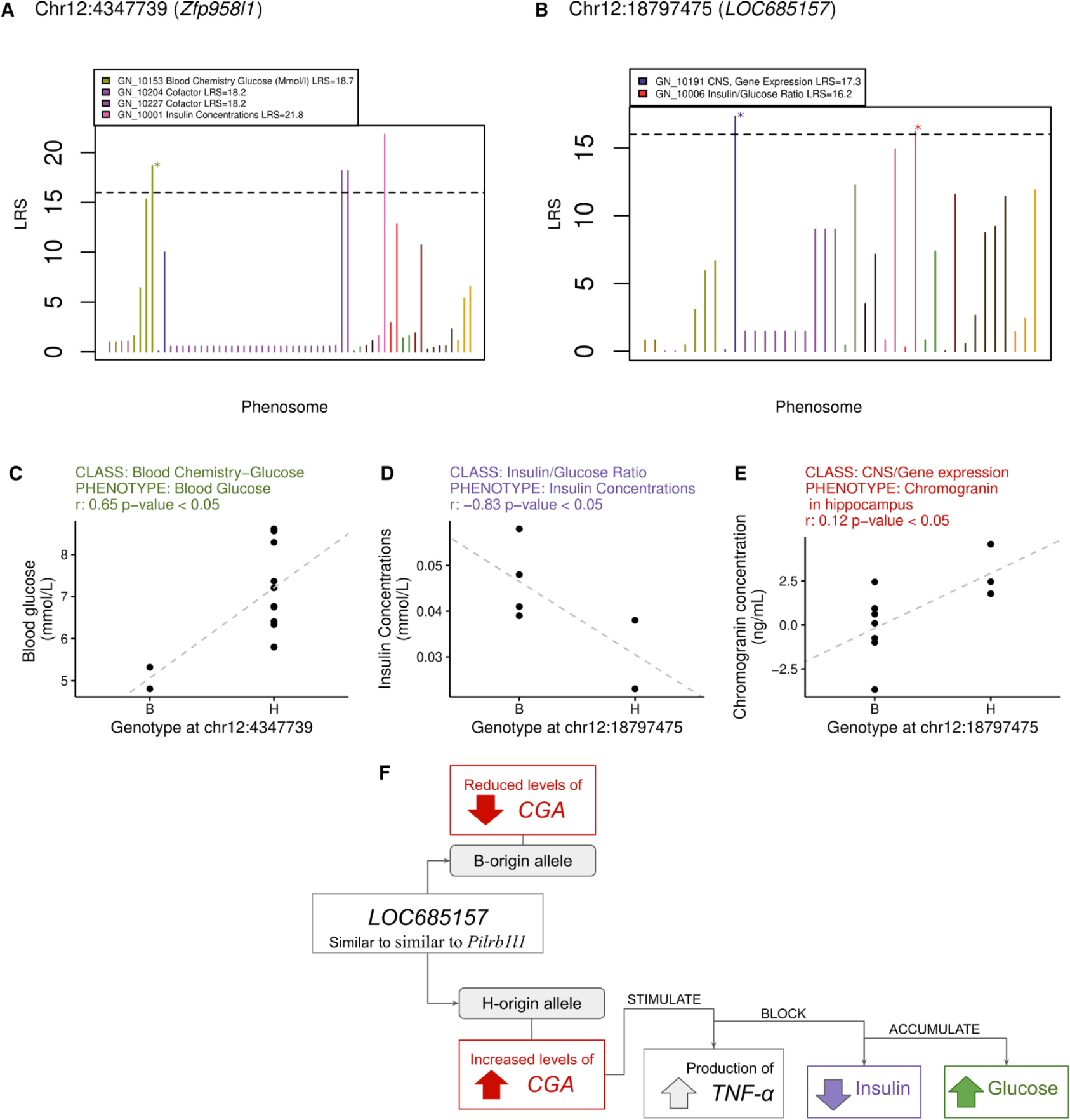
Mapping of validated pangenomic variants against phenotypes in GeneNetwork. **(A-B)** PheWAS and results for two variants in the long non-coding RNAs gene *Zfp958l1* and the annotated gene *LOC685157*, which is similar to *Pilrb1l1*. Colors refer to phenotype classes, the dashed horizontal line indicates threshold for significance in the PheWAS. Stars mark genotype-phenotype associations that were validated also through Spearman correlation analysis (adjusted p-value < 0.05) reported in **(C-E).** In **F** we propose a hypothesis that reconciles our genotype-specific findings with what is known in literature about these genes and their relationship.

The first association was between blood glucose concentration and a SNP (chr12_4347739) within the long non-coding RNAs gene *Zfp958l1*. Rats carrying the *H allele* had significantly higher blood glucose concentrations (**Figure 2C**). chr12_4347739 variant was located within a mapped QTL that controls insulin/glucose ratio, mapped in an independent cohort of 185 F2 rats (Insglur6; LOD score 18.97, p-value: 0.001).^19^

The second relevant association was between a variant and two phenotypes, insulin concentration in blood and gene expression of chromogranin (CGA) in the hippocampus, and an intronic SNP (chr12_18797475) within an annotated locus (*LOC685157*). Rats with the *H allele* at chr12_18797475 had higher expression of CGA in the hippocampus and lower blood insulin concentration (**Figure 2D-E**). The relationship between increased CGA expression, and insulin concentration aligns with previous findings, chromogranin A regulates catecholamine storage and release, variation affecting CGA levels could impact catecholamines that can suppress insulin secretion and affect glucose metabolism.^22,23^ CGA expression is ubiquitous throughout the central nervous system,^20^ especially in the hypothalamus and amygdala regions, and stimulates the production of tumor necrosis factor alpha (TNF-α)^21^ a key inflammatory molecule that promotes insulin resistance and glucose accumulation. We hypothesize that in rats with the *H allele*, increased CGA expression and subsequent TNF-α production leads to insulin resistance, impairing cellular glucose uptake and resulting in reduced blood insulin levels (**Figure 2F**). This proposed mechanism is consistent with the observed reduction in insulin concentration associated with the other significant PheWAS variant (**Figure 2C**).

Remarkably, LOC685157 is similar to the paired immunoglobin-like type 2 receptor beta 1 like 1 gene (*Pilrb1l1,* Ensembl: *LOC681182*) predicted to code for a protein putatively located on the membrane. The associated intronic variant chr12_18797475 was located just 0.7kb upstream of *Pilrb1l1*. *Pilrb1l1* orthologs to human paired immunoglobin like type 2 receptor alpha (*PILRa*) code for a lectin that recognizes both O-glycan and aglycon^20^ and a missense variant (rs1859788, p.Arg78Gly) in the functional immunoglobulin like domain seems to be protective for Alzheimer’s disease.^21^ In mice, *Pilrb1l1* is abundantly expressed on myeloid cells, and required for cell movement out of the circulatory system and towards the site of tissue damage or infection.^22^ *Pilrb−/−* mice have reduced serum or bronchoalveolar lavage fluid levels of tumor necrosis factor alpha (*TNF-α*)^23^ suggesting that increased CGA expression and consequent *TNF-α* production might compensate for the lack of function of *Pilrb1l1*. Finally, chr12_18797475 is located 0.8 Mb from a mapped QTL that controls hyperglycemia related to diabetes and increased susceptibility to autoimmune disorder (Iddm2; LOD 18.97).^19^

### Novel complex structural variants discovered from the graph pangenome

SVs (>50bp in length) of chr12 from 31 HXB were called using four assembly-based methods (pav, svim, hall, vg)^8^, one of which was a graph-based method (vg). We retained 7,811 SVs supported by at least two methods, out of 9,405 identified by at least one method **(Figure S3)**. Contrary to small variants, here the graph-based method called less SVs than other methods (**Figure 1E**). Out of the 7,811 SVs, 1,282 were insertions and 6,526 deletions. We predicted the functional consequences of variants on genes and proteins **(<u>Table S8</u>, Figure 1F-G**). The majority of variants were classified as modifiers (56.0%). A big proportion of variants had a high impact (44%) and were located within 918 protein-coding genes. Notably, the majority of variants were located in exon regions and introns **(<u>Table S8</u>)**.

We further validated SVs of the SHR/OlaIpcv sample called from the pangenome. We retained 2,481 variants supported by at least two methods out of 4,853 identified by at least one method **(Figure S3)**. We selected ten variants with high impact and sized between 50-500bp, that were examined using Oxford Nanopore long-read data from SHR/OlaIpcv rat. Two insertions and four deletions were confirmed, suggesting a validation rate of 0.6 **(Figure S4)**.

All six validated SVs were detected in additional non-SHR strains, with variant frequency varying from 0.35 to 1 **(<u>Table S9</u>)**. A deletion of 300bp in the Lmtk2 gene was found in SHR/OlaIpcv and all other samples (**Figure 3A**). A second deletion of 357bp in the *Mcemp1* gene found in SHR/OlaIpcv was also found in 17 rats (**Figure 3B**). This G-rich SV contained a non-LTR retrotransposon that belongs to the Long Interspersed Elements (LINEs). It also contained two B1F repetitive elements, or *Alu,* that belong to the class of retroelements called short interspersed elements (SINEs, **<u>Table S9</u>**).

**Figure 3.**
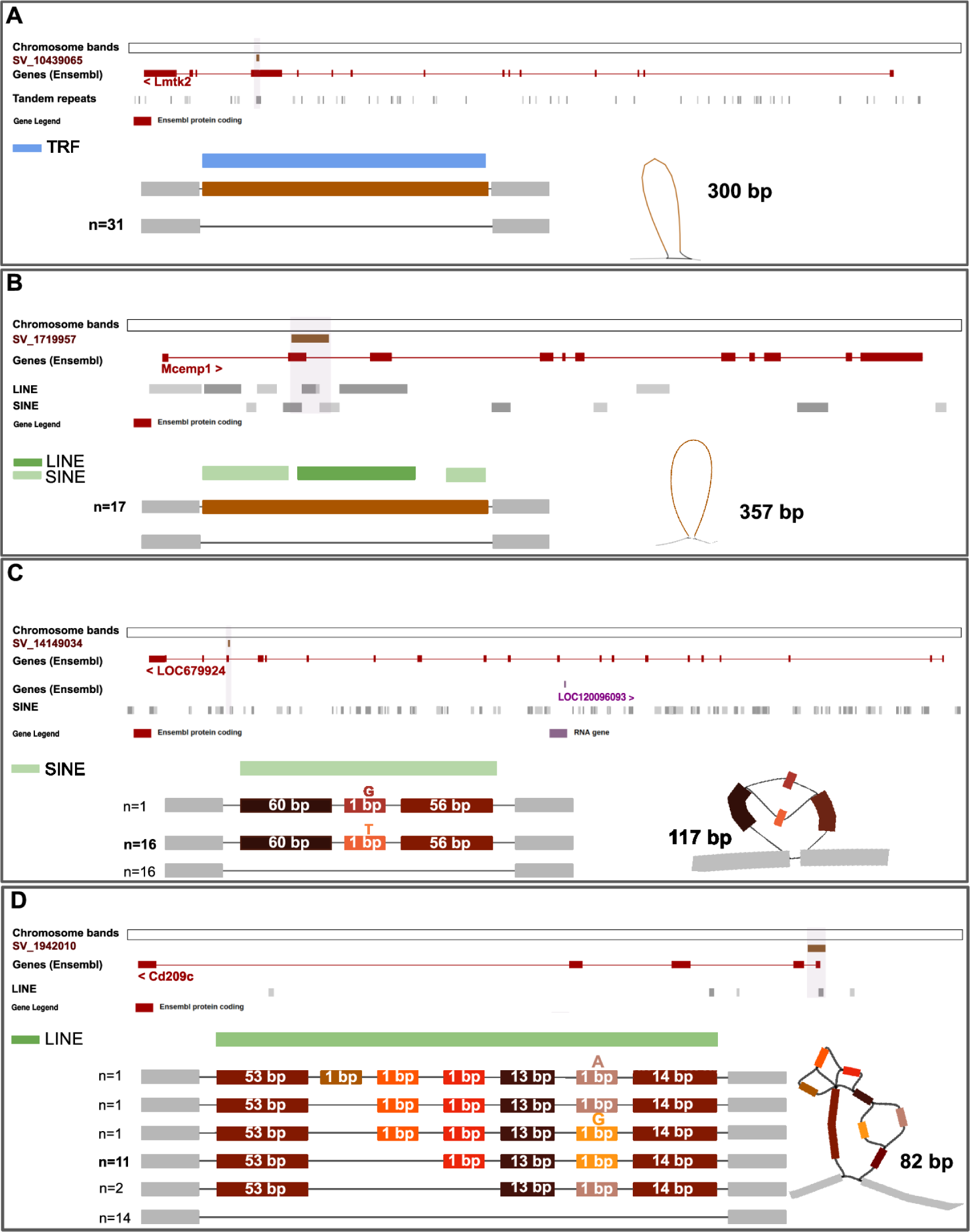
Resolution of complex haplotypes in validated structural variants (SVs) discovered using pangenome graphs. Visualization using ODGI and Bandage of four SVs predicted to have high impact and validated with Oxford Nanopore sequencing technology. **(A)** A deletion of 300bp in the *Lmtk2* gene supported by all four methods of identification, is found in all 31 samples and contains a Tandem Repeats element (TRF). **(B)** A deletion of 357bp in the *Mcemp1* gene is supported by all four methods and is found in 17 rats. The gene contains SINE and LINE elements. **(C)** A deletion of 117 bp in the *LOC679924* gene is supported by two methods, and includes a single nucleotide variant. The SV overlaps for 80 bp with a SINE element. **(D)** A complex insertion of 82 bp in the *Cd209c* gene, supported by all four methods. The insertion resolves in six haplotypes of which two are most abundant, and falls completely in a non-LTR retrotransposon (LINE).

One deletion and one insertion found in SHR/OlaIpcv represent examples of complex variation that were fully resolved by the pangenome graph that captured all the realized haplotypes. The first complex deletion SV (117 bp, supported by two methods) was found in the *LOC679924* gene, predicted to enable ATP binding activity, and overlapped for 80bp with a SINE element (**Figure 3C**). The second complex insertion SV (82 bp, supported by all four methods) fell within the *Cd209c* gene, and, despite its shorter length, was made of seven variable blocks, which combined into six haplotypes, four of which were seen less than two times (**Figure 3D**). It contained a Lx8b_3end, a non-LTR retrotransposon in rodents (LINE). The Human ortholog of *Cd209c* is implicated in Coronavirus infectious disease,^24^ aspergillosis,^25^ dengue disease,^26^ and tuberculosis.^27^

## Discussion

In this study, we built the first rat pangenome to explore the genetic variation landscape of the rat HXB/BXH family. We conduct association mapping with emphasis on novel variants identified by our pangenomic approaches on chr12. The pangenome graph we built enabled the discovery of novel small variants and SVs. Although the sequencing data we used (i.e., 10x Linked-Reads) were not ideal, our research highlights the potential for adequate pangenomic reconstruction. One key finding of this study lies in the significant increase in the number and complexity of variants identified by the pangenome.

After removing uncertain sequence stretches (Ns) the ratio between pangenome and reference sequence lengths is 1.05. Variants in the same genomic regions across multiple samples lead to multiple representations of that region, and the graph accounts for all the different alleles. For this reason, pangenome graphs can be longer than the individual genomes they represent. In particular, we observed in the pangenome 0.18 Gb of non-reference sequence with respect to mRatBN7.2. This figure become 3.9 Gb when including Ns: such high value overestimate the real non-reference bases in our pangenome and reflect the technological limitation of applying Linked-Reads in *de novo* assembly. Most of the novel pangenomic variation is complex, confirming that the pangenome captures genetic variation beyond that contained in the reference genome.

Validation in SHR/OlaIpcv shows high rates of concordance and, except for complex repeated regions, we validated through resequencing novel variants in challenging regions. We provide validated examples of how the pangenome fully resolves complex nested variants within repetitive regions. We identified and validated novel complex SVs falling within genes implicated in immunity and enriched for transposable elements, a major source of genetic polymorphism, known to act as regulatory elements within immune regulatory networks, and facilitate rapid adaptive evolution of immune responses.^28^ Mutations in the *Lmtk2* gene have been linked to multiple neurological disorders, including Alzheimer’s disease, Parkinson’s disease, and schizophrenia in humans. *Mcemp1* seems to be involved in the regulation of mast cell differentiation and immune responses. Its human orthologous *MCEMP1* is one of the top inducible genes in many inflammatory diseases, such as asthma,^29^ idiopathic pulmonary fibrosis,^30^ cancer,^31^ sepsis,^32^ and stroke.^33^ *MCEMP1* is a critical factor in allergic and inflammatory lung diseases,^34^ with the level of *MCEMP1* expression positively correlating with disease severity. These examples provide crucial insights into regions of the genome that play a key role in adaptation, but have been poorly investigated so far due to their challenging resolution.

We demonstrate that novel variants identified by the pangenome are valuable for mapping quantitative traits, demonstrating that access to this fullest spectrum of genetic diversity is key for studying genotype-phenotype connections. We used validated SNPs to conduct phenome-wide association analysis, to discover two significant associations: blood glucose concentration linked to a variant (chr12_4347739) within the *Zfp958l1* gene, and insulin levels in the blood and the expression of chromogranin in the hippocampus, mediated by an intronic variant (chr12_18797475) within a gene (*LOC685157*) similar to *Pilrb1l1*.

Overall, this pangenomic study enhances our understanding of the rat HXB/BXH family and sets a new standard for genomic research in model organisms, underscoring the importance of comprehensive genetic analysis for revealing the complexity of genotype-phenotype connections.

### Limitations of the study

Although this pioneering rat pangenome analysis demonstrates the capability to capture complex variation beyond the reference genome and enable more complete genotype-phenotype mapping, we note a few limitations. The Linked-Reads technology is not ideal for *de novo* assembly and leads to an overestimation of non-reference sequences due to unresolved regions. Additionally, complex regions remain challenging to fully resolve despite validation efforts. The combination of Linked-Read and long-read sequencing technologies offers the promise of constructing higher-quality assemblies and then pangenomes. As sequencing technologies and algorithms continue advancing, we expect to achieve complete pangenome assembly and whole genome structural variation characterization that will reveal the fullest extent of genome variation across rat strains. Despite current limitations, our study demonstrates the capability of pangenomes to enable more accurate, even more complete genotype-phenotype mapping.

## Acknowledgments

V.C. was supported by Ministero dell’Istruzione, dell’Universita’ e della Ricerca of the Italian Republic (PRIN 2020J84FAM). The work of H.C. E.G. and R.W.W. is supported by NIH NIDA U01DA047638. M.P. was supported by a program from the National Institute for Research of Metabolic and Cardiovascular Diseases (Program EXCELES, ID Project No. LX22NPO5104) funded by the European Union - Next Generation EU and by grant LUAUS23095 within the INTER-EXCELLENCE program of the Ministry of Education, Youth, and Sports of the Czech Republic.

## Author contributions

Conceptualization, F.V., A.G., M.E., E.G., R.W.W., H.C., V.C.; Formal Analysis, F.V., A.G., M.E., H.E., V.C.; Visualization, E.G., R.W.W., H.C., V.C.; Investigation, R.W., T.G., M.P.; Resources, P.P., R.W.W., H.C.; Data Curation, F.V., A.G., R.W., T.G., E.G., R.W.W., H.C., V.C.; Writing - Review & Editing, F.V., A.G., R.W., T.G., E.G., P.P., R.W.W., H.C., V.C.; Supervision, E.G., R.W.W., H.C., V.C.

## Methods

### Supernova assembly

Fastq files were used as the input for the Supernova (version 2.1.1) program provided by 10x genomics^11,12^ to generate *de novo* assembly with default settings. Complete information about the depth per sample can be found in ^10^. To evaluate genome assembly completeness, we used the Compleasm^35^ tool to assess representation of universal single-copy orthologs expected to be present across mammals (BUSCO Mammalia gene set). Compleasm searched each Supernova genome assembly for the presence and completeness of these conserved orthologs. The reference mRatBN7.2/rn7 genome was analyzed comparably as a benchmark.^10^

### Pangenome generation and evaluation

Supernova haploid assemblies obtained by 10x chromium Linked-Reads technology were mapped against the mRatBN7.2/rn7.fa reference genome using wfmash^36^ to partition the assemblies’ contigs by chromosome. Mapped assemblies were used to build the pangenome with PGGB (v.c1c3a1565604fc41f880bccd9f46d0a709f3e774)^37^ using this combination of parameters: -p 98 -s 2000 -n 32 -F 0.001 -k 79 -P asm5 -O 0.03 -G 4001,4507 -V rn7:# -t 48 -T 48. Validation of the call set was performed on SHR/OlaIpcv sample, using one true positive set obtained from joint-calling conducted using GLnexus.^14^ Prior to comparison, the pangenome-derived and the validated call set were processed to remove missing data, sites where alleles are stretches of Ns, homozygous reference genotypes and variants greater than 50bp before normalization and decomposition using bcftools^38^ under default parameters. While the pangenome-derived VCF was based on haploid assemblies, for comparison purposes the calls were considered as homozygous diploid in the assumption that SHR/OlaIpcv is isogenic. Comparison of the two call sets was performed with RTG tools (v.3.12.1)^39^ using the --squash-ploidy option. The RTG tool used for the variants’ evaluation gives us some information about the variants in common, called by vg and JC. From these results we considered only the vcf file which contains variants called by vg-only. RepeatMasker^38^ was used to mask complex regions, easy regions that do not contain low complexity regions and repeats. Hard regions contain the whole genome regions. RepeatMasker, Ensembl Genome Browser^40^ and Dfam^41^ databases are used to check the abundance of repeats across the genome. We mapped raw reads of SHR/OlaIpcv against the pangenome using vg giraffe (v. 1.41.0).^17^ For the pangenome graph statistics we used ODGI (v.0.7.3),^42^ we report the non-reference sequence computed by summing the length of the graph nodes that are not traversed by the reference genome. This way we count each non-reference node only once. For the pangenome graph visualization we used ODGI and Bandage.^43^ To check the length of the rat genome assemblies (mRatBN7.2/rn7 reference) we used the National Center for Biotechnology Information (NCBI) online resource.^44^ For the vcf statistics we used bcftools^38^ stats and vt peek.^45^

### Variant calling

The variant calling of variants on the pangenome was obtained using vg (v. 1.41.0)^15^ and in particular -V rn7:# into the pggb command line. Small variants are variants with length <50 bp, simple variants are SNPs/MNPs/Indels, complex variants are allelic variants that overlap but do not cover the same range. For the variant classification in simple and complex variants we used vt resource (https://genome.sph.umich.edu/wiki/Variant_classification). For variant calling based on the reference genome, fastq files of 10x chromium linked data were mapped against mRatBN7.2 using LongRanger (version 2.2.2) (https://github.com/10XGenomics/longrangere Long Ranger code is available at https://github.com/10XGenomics/longranger). DeepVariant (ver 1.0.0)^38^ was used to call variants for each sample. Joint calling of variants for all the samples was conducted using GLnexus.^14^

To optimize a pangenomic perspective and capture SVs that might be missed with traditional reference-based methods, we applied several SV calling pipelines. As a graph-based method, we used PGGB and vg to generate a comprehensive VCF file. Variants were normalized using vcfwawe^30^ and then filtered by keeping only variants >50 bp. As assembly-based methods, we applied 3 pipelines as described in.^8^ To evaluate the concordance between all the approaches, we conducted an overlap analysis using truvari (v.4.0),^31^ a tool for assessing SV calls following some criteria 1) we merged SVs using a maximum allowed distance of 1000 bp, as measured pairwise between breakpoints (begin1 vs begin2, end1 vs end2) 2) we reported calls supported by at least 2 callers and they have to agree on the type and on the strand of the SV 3) we removed variants with more than 10% of Ns in REF/ALT. For the SV’s pangenome graph extraction we used odgi (v.0.7.3) and for the visualization we used Bandage and odgi. Subsequently, we investigated the impact of small variants and SVs using SnpEff (v5.1),^30^ a well established tool for annotating and predicting variant effects. Through SnpEff analysis, we gained insights into the potential functional consequences of the detected genetic variants on the identified genes.

### Sanger resequencing for variant validation

Variants only found with the pangenome approach were randomly selected from Chr12 for the SHR/OlaIpcv sample. Primers were designed using NCBI primerblast and synthesized by IDT at 25 nm scale with standard desalt purification (Coralville, Iowa). DNA from a male adult SHR/OlaIpcv rat was extracted from brain tissue using DNEasy Blood and Tissue Kit (Qiagen, Cat no. 69504) and amplified using Phusion High-Fidelity PCR Kit (ThermoFisher, Cat no. F553L) in a Thermocycler for 30 cycles. PCR products were inspected on 1% agarose gel. PCR products with one distinctive band were submitted to GeneWiz (South Plainfield, NJ) for sequencing. Sequencing results were mapped against the target sequence using the blast2seq application and mutations were manually examined for confirmation.

### PheWAS analyses and functional annotation

A Phenotype Wide Association Study (PheWAS) was conducted to identify potential traits related to the validated SNPs. The Phenome of the HXB/BXH family was downloaded from GeneNetwork.org (version 2) using an API. Out of 324 phenotypes we included 60 phenotypes, with data from at least 2 observations per phenotype within phenotypic class. Phenotypes were correlated via the ‘hxb.phenotypes’ object, using an adaptation of BXDtools^49^, an R package containing various genetic tools to work in particular with model organisms. BXDtools employs a simpler linear model. Spearman correlations between genotype and phenotype were carried out using R, with p-values adjusted (Bonferroni) according to the number of samples each phenotype contains. The statistical significance level was set to p-adj < 0.05. This relationship was visualized in a genotype-phenotype plot. Ensembl Genome Browser^35^ and Rat Genome Database (RGD)^30^ were used to identify genes. Genome assembly (mRatBN7.2/rn7) for *Rattus norvegicus* was used.^34,4^ To enhance the visualization and understand the distribution of these genes and their variant impacts on chr12, we employed a specialized R package called karyotypeR.^50^

### SVs validation with nanopore adaptive sequencing

To validate the SVs identified, we employed Adaptive Sampling, a nanopore sequencing-specific software-controlled enrichment method. For Oxford Nanopore Adaptive Sequencing, 4.4 µg of DNA was extracted from 22 mg of microdissected rat brain tissue using the Qiagen DNeasy Blood & Tissue Kit and quantified using Nanodrop. The extracted DNA was fragmented using Covaris’ g-TUBE for 30 seconds at 9100 x g at room temperature, for a median fragment size of ∼3kb. Library prep was performed with ONT’s Ligation Kit and libraries were quantified with Qubit. Following ONT’s guidelines for the use of their Ligation Kit with Adaptive Sampling and ONT Community input, 100 fmol of library was loaded onto a MinION flow cell and run for 72 hours with Adaptive Sampling selected in run setup. Nanopore validated reads are mapped against the mRatBN7.2/rn7.fa reference genome with minimap2,^51^ and the bam files are visualized using Integrative Genomics Viewer (IGV).^52^

## Data and code availability

- Linked-read WGS data for the HXB/BXH family is available from NIH SRA. The SRA identifications for each sample is provided in **(<u>Table S10</u>)**.
- All original code has been deposited at GitHub (https://github.com/Flavia95/HXB_rat_pangenome_manuscript/).

## Supplementary Information

**Figure S1.**
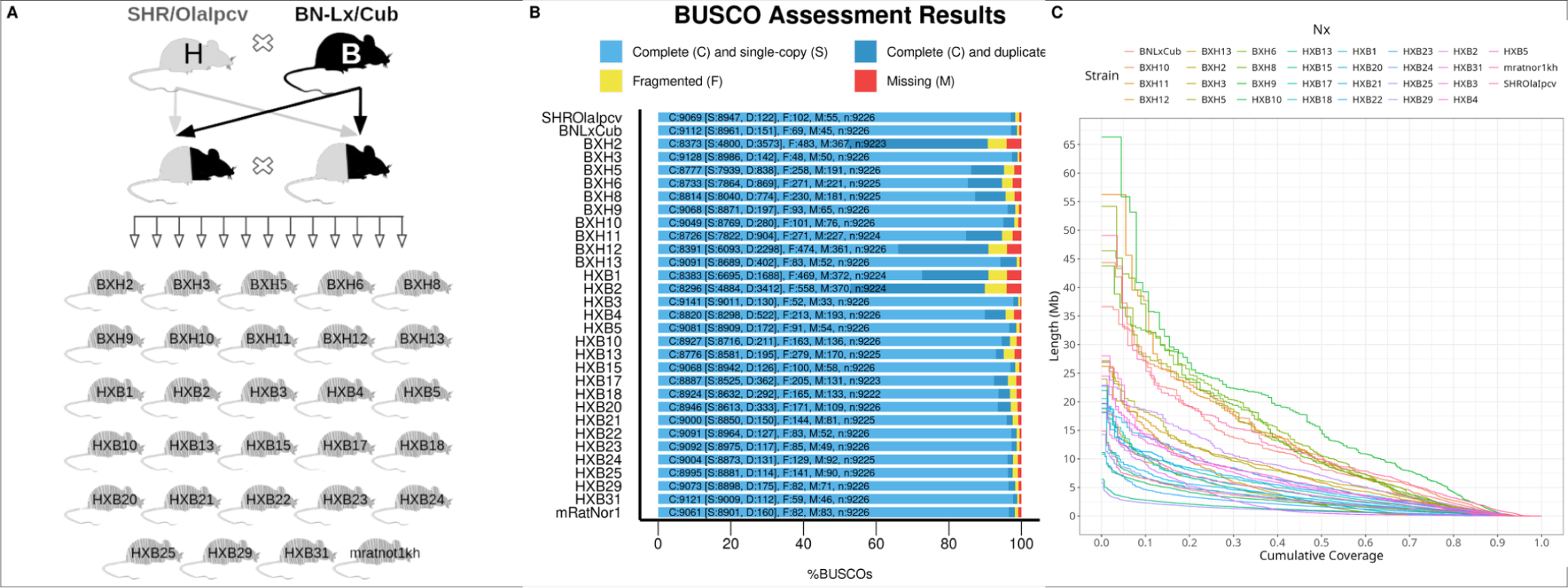
Description of rats used in this study and quality of the genome assembly. **(A)** Overview of the HXB/BXH recombinant inbred rat family describing the origins and breeding history of the HXB/BXH rats. **(B)** BUSCO completeness of the genome assembly used to build the pangenome for genomics data quality control. Bar charts show proportions classified as complete (C, blues), complete single-copy (S, light blue), complete duplicated (D, dark blue), fragmented (F, yellow), and missing (M, red). **(C)** Nx plot showing assembly contiguity for each of the 31 rat strains.

**Figure S2.**
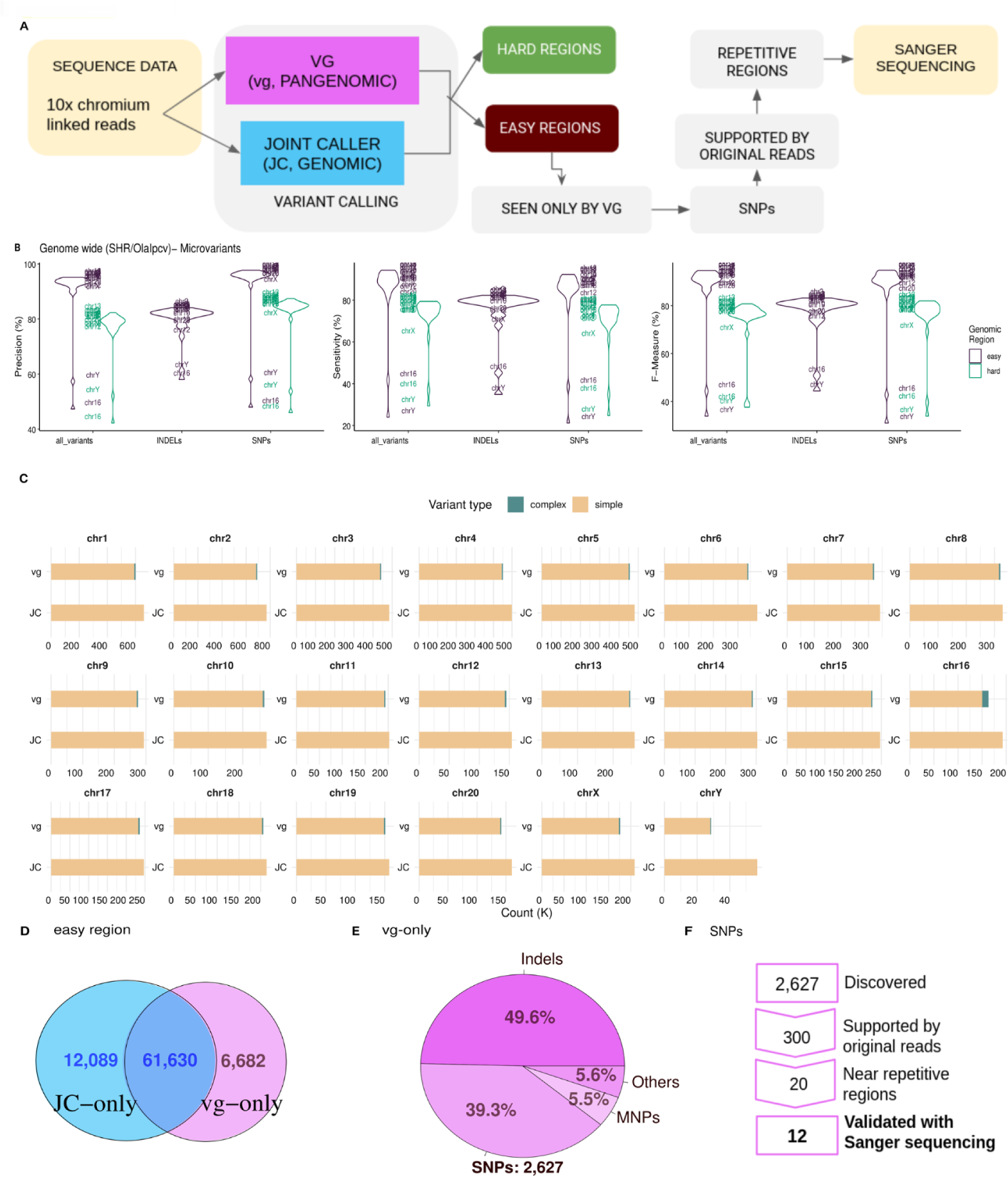
Validation in the SHR/OlaIpcv sample of the small variants called from the genome graph using vg. Small variants were validated over the joint-calling (JC) call set (gold standard) in the SHR/OlaIpcv sample according to the scheme in **(A). (B)** Genome-wide accuracy of vg calls is ∼100% in the easy regions for Single Nucleotide Polymorphisms (SNPs), ∼90% in the easy regions and ∼80% in the easy regions for Indels and hard regions of the genome. Exceptions are seen for chromosomes 16 and Y, which are enriched for complex variation **(C)**. Validation through Sanger sequencing was restricted to easy regions **(D)**, to SNPs **(E)** supported by original reads and in challenging, repetitive regions **(F).**

**Figure S3.**
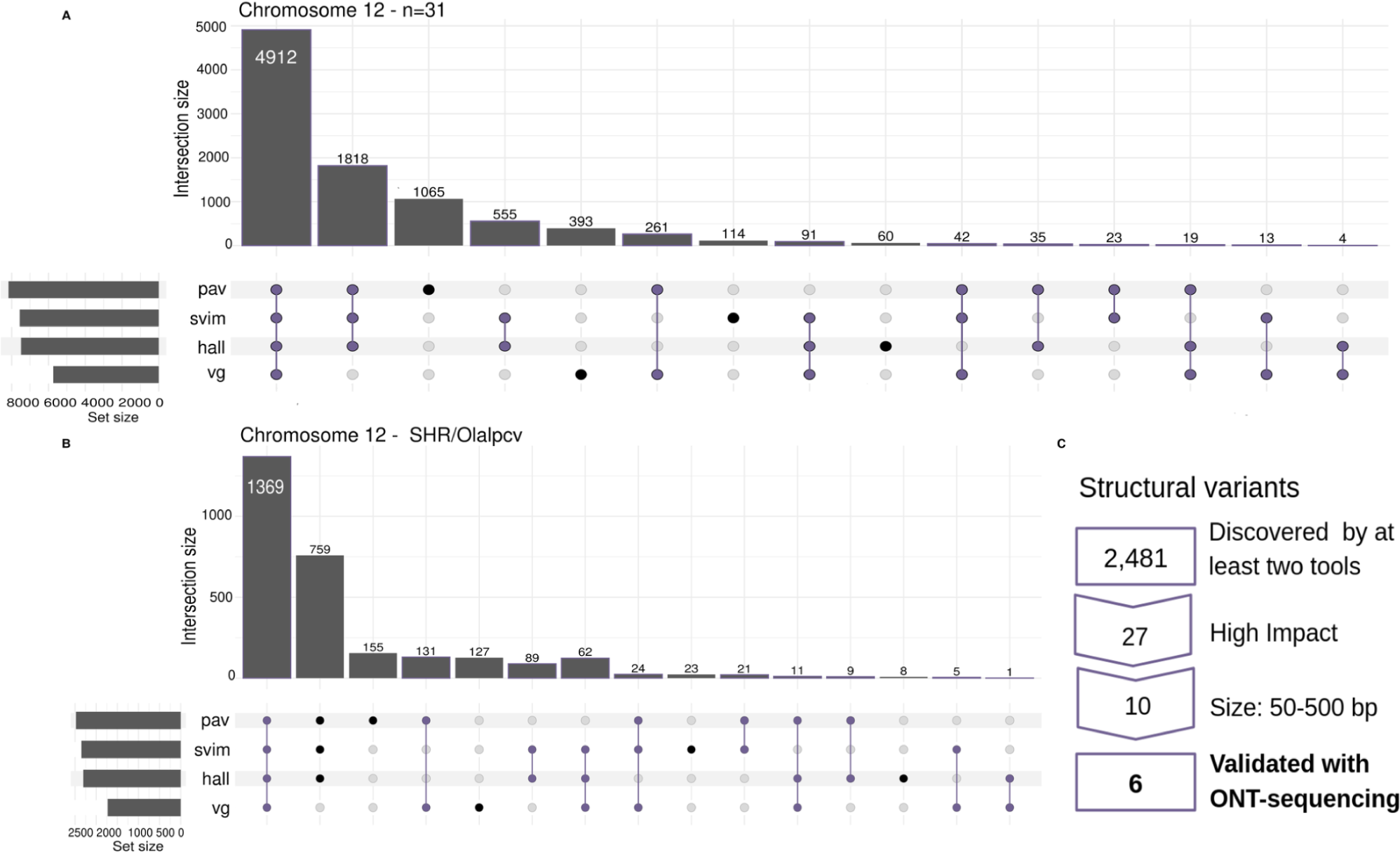
Validation in the SHR/OlaIpcv sample of the Structural Variants (SVs) called from the genome graph using vg. **(A)** Overlap of the call sets obtained by the three assembly-based methods (pav, svim, hall), and the graph-based method (vg) for chromosome 12 using the data from all rats. **(B)** Same as (A) for SHR/Olalpcv only. **(C)** Scheme of the validation for SVs

**Figure S4.**
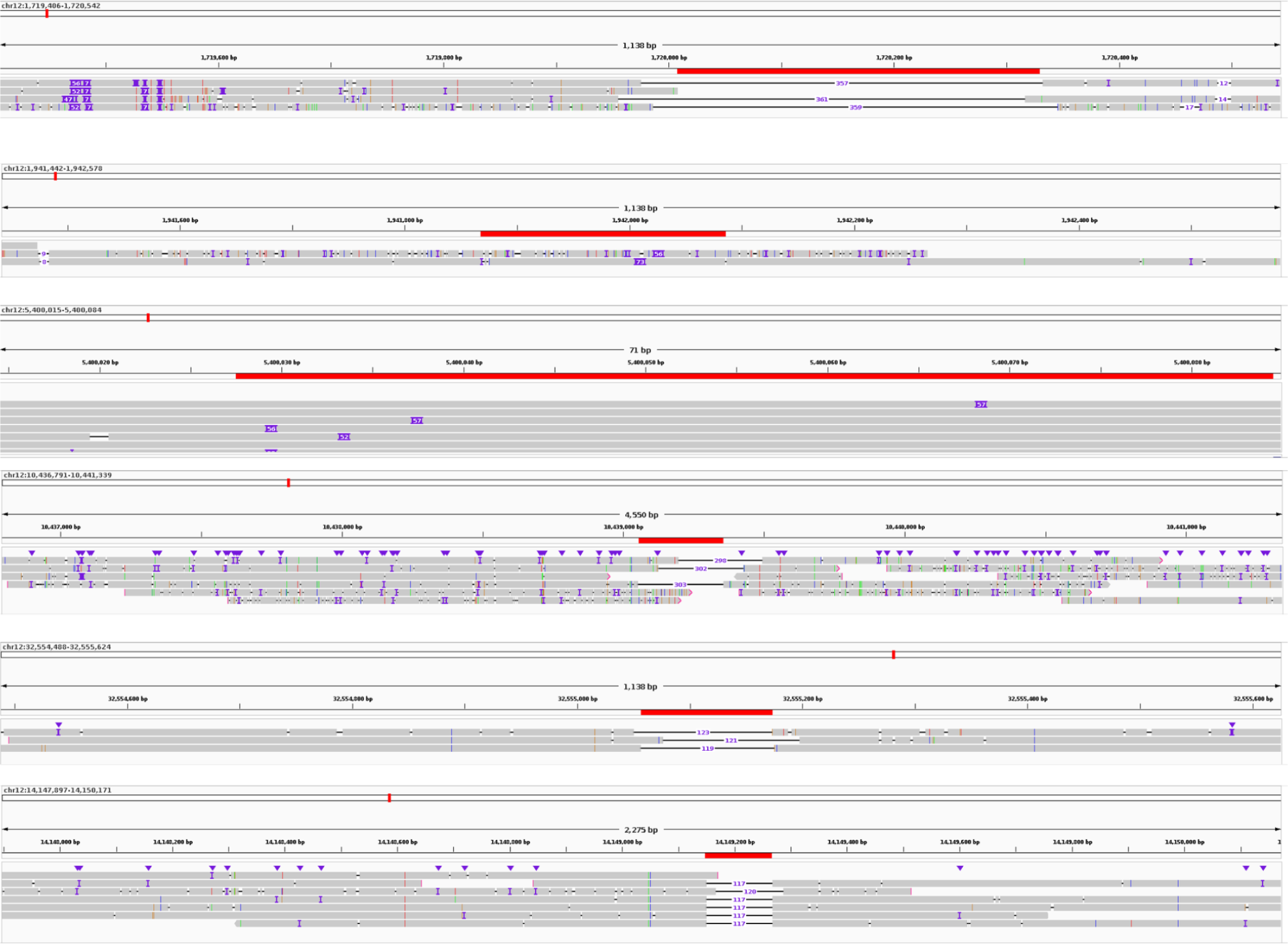
Integrated Genomic View of the validated Structural Variations (SVs) For each SV gray bars are validated reads mapped against the mRatBN7.2/rn7.fa reference genome, red bars define the boundaries of the SV.

**Tables:**

https://docs.google.com/document/d/1qXAEqRID3faaI3BWApNtz_94jV1LXLF4QntjmzwMcrY/edit?usp=sharing

